# Human interferon stimulated genes target ancient features of animal and bacterial viral replication

**DOI:** 10.64898/2026.06.11.731453

**Authors:** Samantha G. Fernandez, Jill E. Hutchinson, Joel M.J. Tan, Sonomi Yamaguchi, Anne A. Roffler, Aaron G. Schmidt, Philip J. Kranzusch

## Abstract

Animal and bacterial cells defend against viral infection by rapidly activating antiviral restriction factors. In human cells, antiviral immunity is initiated by interferon signaling that results in expression of hundreds of interferon-stimulated genes (ISGs)^1,2,3,4,5,6^. Complex regulatory networks and co-evolution of viral evasion strategies complicate analysis of immune proteins under native conditions and the function of most individual ISGs remains unknown^7,8,9,10,11,12^. Here we discover that heterologous expression of ISGs in bacteria is sufficient to protect against infection by diverse bacteriophages demonstrating properties of antiviral restriction preserved across billions of years of viral evolution. A screen of 306 human ISGs against 11 *E. coli* phages reveals that ISGs can restrict phage replication with potency equal to endogenous bacterial defense systems. We select for phage mutants that escape ISG restriction and identify the phage DNA primase-helicase complex as a target of human SPSB1. A 2.0 Å crystal structure of the SPSB1–primase complex uncovers a recognition mechanism of foreign DxNxN protein motifs found in many animal and bacterial viral replication proteins. We show that SPSB1 in human cells recognizes and induces degradation of protein targets containing this motif from norovirus and poxvirus pathogens. Our results establish a cross-kingdom approach to studying immune function and reveal that human ISGs target features of viral replication common across kingdoms of life.

## Introduction

The mammalian interferon response induces expression of hundreds of immune proteins that together drive resistance to viral infection^1,13,2,5,6^. Most ISGs function through unknown mechanisms^11^, limiting molecular analysis of human and animal antiviral immunity. Motivated by recent experiments demonstrating a shared evolutionary connection between human antiviral immunity and bacterial anti-phage defense^14,15,17,18,19,20,21,22,23,24,25,26,27^, we reasoned that experimental models of phage infection may provide a unique opportunity to measure the ability of ISGs to inhibit viruses that have not previously encountered human defenses and reveal molecular strategies human cells use to combat infection. We constructed a synthetic library of 306 human ISGs codon-optimized for bacterial expression and tested each human ISG individually in *E. coli* using an inducible plasmid system previously optimized for analysis of native bacterial anti-phage defense^28,29^ (**Figure 1a**). In a pilot screen with two highly divergent phages T4 and Bas25, we observed that expression of individual human ISG proteins reduced phage titers between 1,000- and 10,000-fold compared to control cells expressing GFP (**Figure 1b**). Replication of phages T4 and Bas25 was restricted by distinct sets of human ISGs, including well-characterized human immune proteins like IFIT1, SLFN5, and MOV10 exhibiting phage-specific phenotypes (**Figure 1b,c**). To control for possible indirect effects on host cell fitness, we measured the impact of human ISG expression on *E. coli* growth and verified that the top hits inhibited phage replication in the absence of cellular toxicity, supporting the robustness of our approach in reconstituting potential *bona fide* human immune protein function in an heterologous bacteria-phage model of infection (**Figure 1d**).

**Figure 1:**
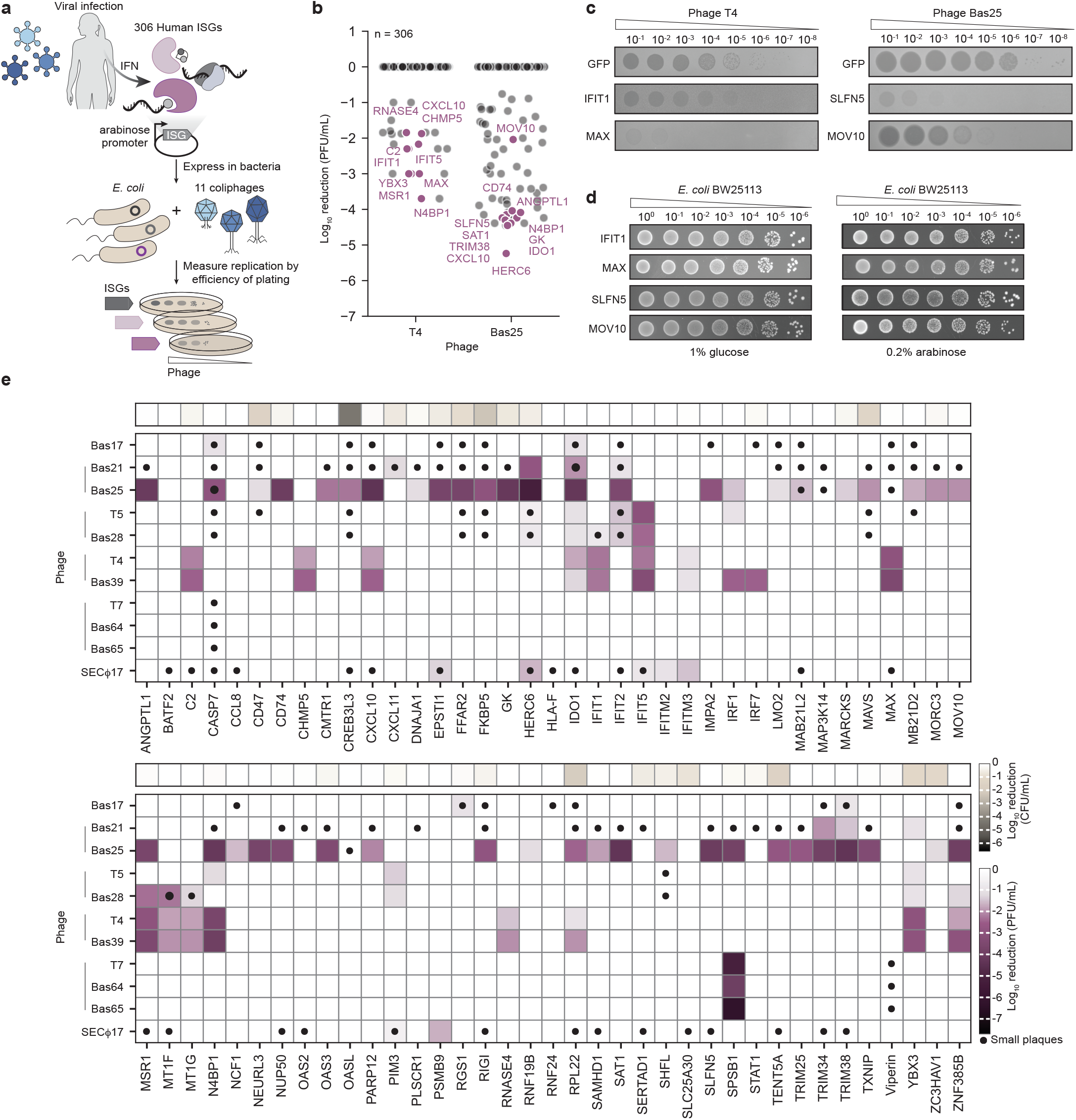
A cross-kingdom screen in bacteria identifies human ISGs capable of restricting phage replication. **a**, Schematic of screen in bacteria to identify human ISGs with anti-phage activity using efficiency of plating (EOP) as a readout. **b**, Viral titer measurements are reported as log_10_ transformed plaque-forming units per mL (PFU/mL) from pilot ISG screens with phages T4 and Bas25. **c**, Representative EOP assays of serially diluted phage T4 and phage Bas25 plated on *E. coli* expressing a GFP control or the indicated human ISG. **d**, Representative toxicity spot assays of serially diluted *E. coli* transformed with the indicated ISG. Samples were plated on LB supplemented with either 1% glucose (promoter off) or 0.2% arabinose (promoter on). **e**, Heatmap of reduction in phage replication (PFU/mL) upon human ISG expression against 11 bacteriophages. Depicted are ISGs with at least a 10-fold-reduction in phage titer or changes in plaque size where color reflects titer measurements relative to GFP control cells. Closed circles indicate small plaque size morphology. Heatmap of reduction in host cell growth in colony forming units per mL (CFU/mL) are aligned at the top of each row. Lines next to phage labels indicate phage family membership (SECΦ17 is a ssDNA phage). Data represent mean of two independent replicates.

We next performed a large-scale screen of human ISG activity against 10 diverse double-stranded phages and 1 single-stranded DNA phage representing 6 viral families. Cross-kingdom analysis revealed 74 human ISGs capable of restricting phage replication, with human ISG-dependent defense phenotypes ranging from changes in plaque morphology to multi-log reductions in viral titer (**Figure 1e and Extended Data Figure 1**). Human ISG-dependent anti-phage activity was largely viral family specific (**Extended Data Figure 2a,b**), with ISGs that caused atypically broad defense phenotypes across families more associated with increased cellular toxicity (**Extended Data Figure 2c,d**). Forty of the top human ISG hits in our screen exhibited significant protection against phage infection with no detectable cost to *E. coli* cellular fitness (**Figure 1d and Extended Data Figures 1 and 2**). Remarkably, the strongest ISG hits in our screen including human RNASE4, SPSB1, and TRIM25 proteins exhibited >100,000–1,000,000-fold reductions in phage titers similar in magnitude to the effects of native *E. coli* bacterial defense systems specifically evolved for inhibition of phage replication^28,20,21,30^ (**Figure 1d and Extended Data Figures 1 and 2**). Together, these results demonstrate that individual human ISGs are sufficient to protect prokaryotic cells from viral infection and establish cross-kingdom analysis as a heterologous model to directly interrogate human immune protein function.

### Analysis of human ISG function in bacterial anti-phage defense

To begin to define the specificity of ISG-mediated immunity, we analyzed defense against phage replication in bacteria compared to previous screens of ISG function in human cells. Consistent with ISG over-expression screens against human viruses^7,8,31,32,33,34,6^, we observed that active ISGs were more likely to restrict replication of closely related phages within the same family compared to unrelated phages (**Figure 2a**). In particular, a set of ISGs including CHMP5 and RNASE4 restricted Myoviridae phages (T4 and Bas39) and a distinct set of ISGs including viperin and SPSB1 restricted Autographiviridae phages (T7, Bas64, Bas65), supporting that ISGs target specific features of replication shared within viral families (**Figure 2a**). We observed broad ISG-mediated immunity against Siphoviridae phages (Bas21 and Bas25), with Bas25 being particularly susceptible and inhibited by 46 ISGs, whereas Autographivridae phages were largely resistant to human ISG function (**Figure 2b**).

**Figure 2:**
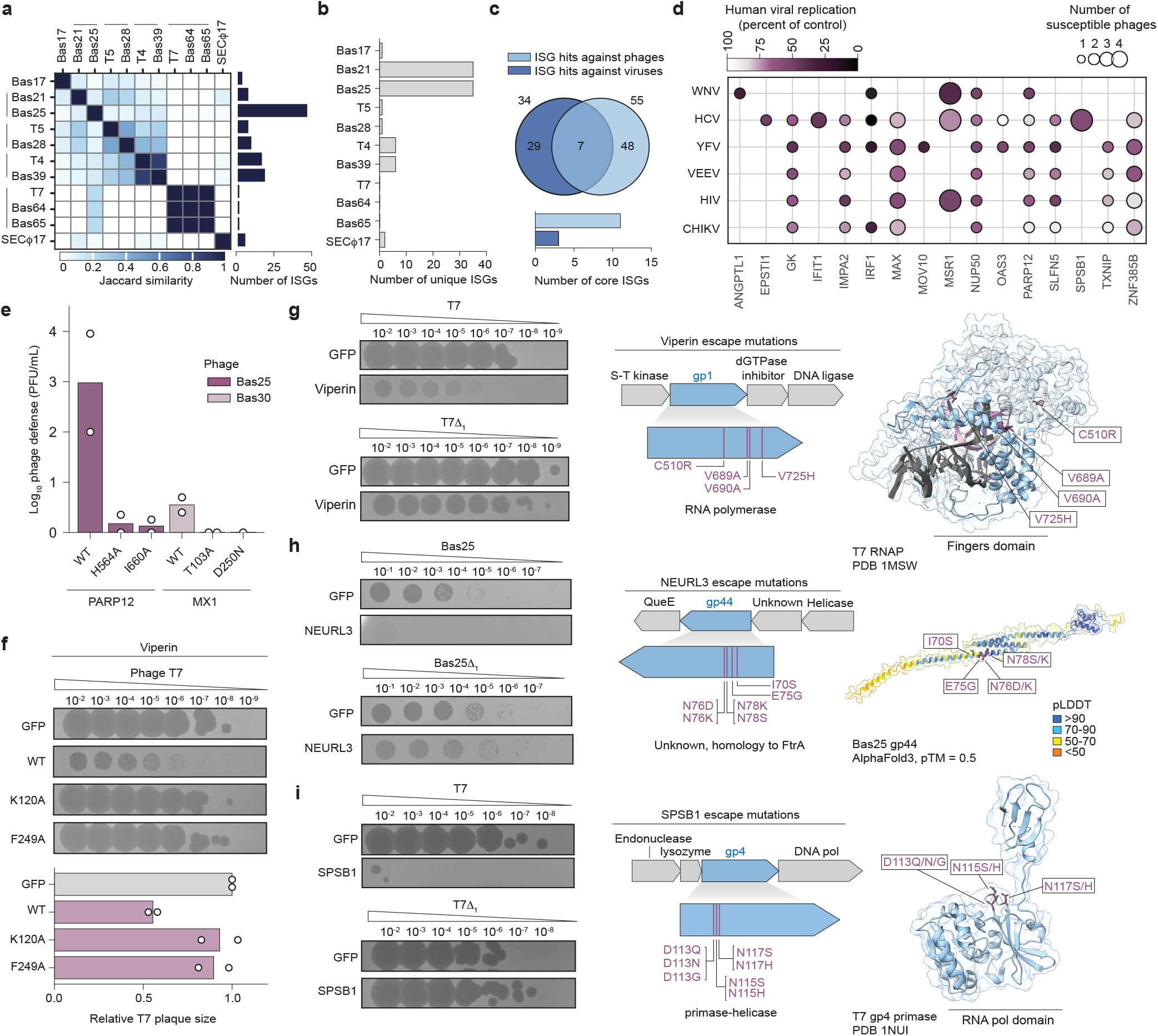
Human ISGs function as immune effectors in bacteria that can induce immune evasion in phages. **a**, Heatmap depicting similarity (Jaccard index) between the sets of ISGs that restrict each phage (left) along with the total number of ISGs that restrict each phage (right). Increasing Jaccard index number indicates increasing similarity. **b**, Number of ISGs that uniquely restrict each family of phages. **c**, Venn diagram showing intersection of top ISG hits in screen against human viruses^7^ (dark blue) and top hits in screen against phages (light blue) (top). Number of ISGs from each respective screen annotated as a “core ISG” (bottom). **d**, Bubble plot depicting the strongest ISG hits in the bacterial phage screen (>100-fold-reduction) and the previously described antiviral activity of each ISG against human viruses^7^. Color depicts strength of antiviral response against animal viruses as a percent of viral titer relative to control cells whereas bubble size indicates number of phages each ISG defended against. **e**, Quantification of EOP assays of indicated ISGs and variants challenged with phages Bas25 or Bas30, n=2. **f**, EOP assays (top) and quantification (bottom) of plaque size depicting dilutions of phage T7 on indicated viperin variants. Plaque size was measured relative to GFP control cells, n=2. **g,h,i**, EOP assays of *E. coli* expressing indicated ISGs against serial dilutions of either the parental phage or phage escape mutant (left). Schematic of mapped phage escape mutants and corresponding genomic loci (middle). Subset of escape mutations mapped onto corresponding structures of each gene identified by escape mutant analysis (right).

Comparison of the strongest ISG hits in our screen (>100-fold reduction in phage replication or significant changes in plaque size) against ISGs previously identified to inhibit human viruses^7^ revealed that seven ISGs including IRF1 and MOV10 defended against both phages in our panel and select human viruses (**Figure 2c,d**). Four ISGs in our screen were previously found to reduce human viral titers by at least 50% while the majority had moderate phenotypes (20–40% reduction in replication) and in some cases had no impact on viral replication against the tested human viruses (**Figure 2d**). Interferon signaling is known to induce expression of a highly variable set of genes in different animal species with a minimal set of 62 “core ISGs” shared across diverse vertebrates^5^. Core ISGs more frequently appeared as hits against phages compared to previous experiments in human cells (**Figure 2c and Extended Data Figure 3a**), highlighting the utility of a heterologous cross-kingdom screen to reveal immune phenotypes by testing viruses that have no prior exposure to animal antiviral defenses.

We sought to validate our results by testing a series of catalytically inactive mutants of ISG hits with well-defined enzymatic functions. Mutations to PARP12 known to inhibit polyADP-ribose polymerase function^35^ and to MX1 known to inhibit GTPase activity^36^ abrogated all ability for these human ISGs to restrict phages Bas25 and Bas30 (**Figure 2e and Extended Data Figure 3c**). Likewise, mutations to the S-adenosylmethionine (SAM) enzyme viperin known to inhibit catalytic activity or SAM cofactor-binding^37^ restored replication and normal plaque morphology of phage T7 agreeing with prior evolutionary studies of viperin proteins in bacterial anti-phage defense^19^ (**Figure 2f**). Finally, we tested mutations in the RNA helicase MOV10 reported to disrupt putative helicase activity^37^ and observed no impact on the phage defense phenotype suggesting that RNA-binding and not helicase activity may be responsible for immune restriction in this heterologous context (**Extended Data Figure 3d**). Together, these results confirm that ISG-mediated immunity in bacteria is overall dependent on canonical protein function.

### Phages rapidly evolve resistance to human ISG immunity

Analysis of mutant phages that escape bacterial immunity has been a key approach to determine the molecular mechanisms controlling many anti-phage defense systems^38,39,40,41^. Having demonstrated that human ISGs can function as potent immune effectors in bacteria, we next screened phages for the ability to spontaneously escape ISG-mediated immunity and successfully isolated mutant phages resistant to human ISGs including viperin, NEURL3, and SPSB1 (**Figure 2g–i and Extended Data Figure 3e**). Viperin restricts replication of human viruses including hepatitis C virus (HCV) by catalyzing synthesis of a modified nucleoside analog ddhCTP (3′-deoxy-3′,4′-didehydro triphosphate nucleotide) that inhibits viral replication complexes and terminates chain extension^42^. We isolated and sequenced 12 escape mutants in the phages T7, Bas64, and Bas65 and observed that all clones resistant to human viperin inhibition had single missense mutations in the viral RNA polymerase gene *gp1* (**Figure 2g**). Mapping 9 unique mutations on the phage T7 RNA polymerase structure^43^ revealed that each mutated residue was located in the polymerase “fingers” domain responsible for incoming nucleotide selection consistent with protein mutations that likely alter substrate selection and limit ddhCTP incorporation. Human NEURL3 (neutralized E3 ubiquitin protein ligase 3) has been shown to restrict HCV replication through an unknown mechanism^44^. We isolated and sequenced 10 escape mutants in phage Bas25 and identified 11 unique mutations with 4 occurring in the same two residues in an uncharacterized viral gene *gp44*. We predicted the structure of gp44 with AlphaFold3^45^ and observed that the escape mutations cluster on a predicted helical interface suggesting the activity of NEURL3 may target the function of this region to restrict phage replication (**Figure 2h and Extended Data Figure 3e**). Finally, SPSB1 (SPRY and SOCS-box domain containing protein 1) is a member of a poorly characterized gene family in humans that has not previously been proposed to have a direct role in viral restriction but has been reported to impact regulation of NF-κB, iNOS, TGF-β signaling in immune cells^46,47,48,49^. We isolated and sequenced 21 escape mutant phages resistant to SPSB1 restriction and identified 7 unique mutations in the phage T7 primase-helicase gene *gp4* (**Figure 2i and Extended Data Figure 3e**). All isolated mutations mapped on the gp4 crystal structure^50^ to three residues D113, N115, and N117 that reside in a flexible loop near the NTP binding site^51^, suggesting that SPSB1 interferes with an essential function of the viral primase-helicase protein. Together, these results reveal that phages can rapidly acquire resistance to human ISG-mediated immunity and demonstrate that bacteria-phage genetic approaches can be directly applied to the study of human immune protein function.

### Structural basis of human SPSB1 inhibition of phage replication

We next focused on defining the mechanism driving human SPSB1 inhibition of phage infection. SPSB1 induces an incredibly potent >1,000,000-fold reduction in replication of Autographiviridae phages T7, Bas64, and Bas65 with no associated cellular toxicity during *E. coli* expression (**Figure 1**). The SPSB1 protein is comprised of a central SPRY domain that in other proteins is typically involved in ligand recognition^52^ and a C-terminal SOCS-box domain that has been shown to recruit the CRL5 E3-ligase complex^46^. Analyzing SPSB1 deletions, we observed that the SOCS-box domain was dispensable and that a construct containing only the SPSB1 SPRY domain was sufficient to protect bacteria against phage T7 infection (**Extended Data Figure 4a**). Because all phage T7 escape mutants mapped to the viral primase-helicase protein gp4 (**Figure 2**), we co-expressed the human SPSB1 SPRY domain and phage T7 gp4 and observed that the proteins copurify and strongly interact to form a stable 1:1 complex on size-exclusion chromatography (**Figure 3a,b and Extended Data Figure 4b**). We observed that complex formation specifically required the gp4 primase domain and used biolayer interferometry (BLI) to determine that SPSB1 binds to the gp4 primase with nanomolar affinity (K_d_ = 126 nM) (**Figure 3c**). Introduction of the phage T7 gp4 escape mutations N115H and N117S disrupted protein interaction *in vitro* demonstrating that these residues are critical for SPBS1–gp4 complex formation (**Figure 3d**).

**Figure 3:**
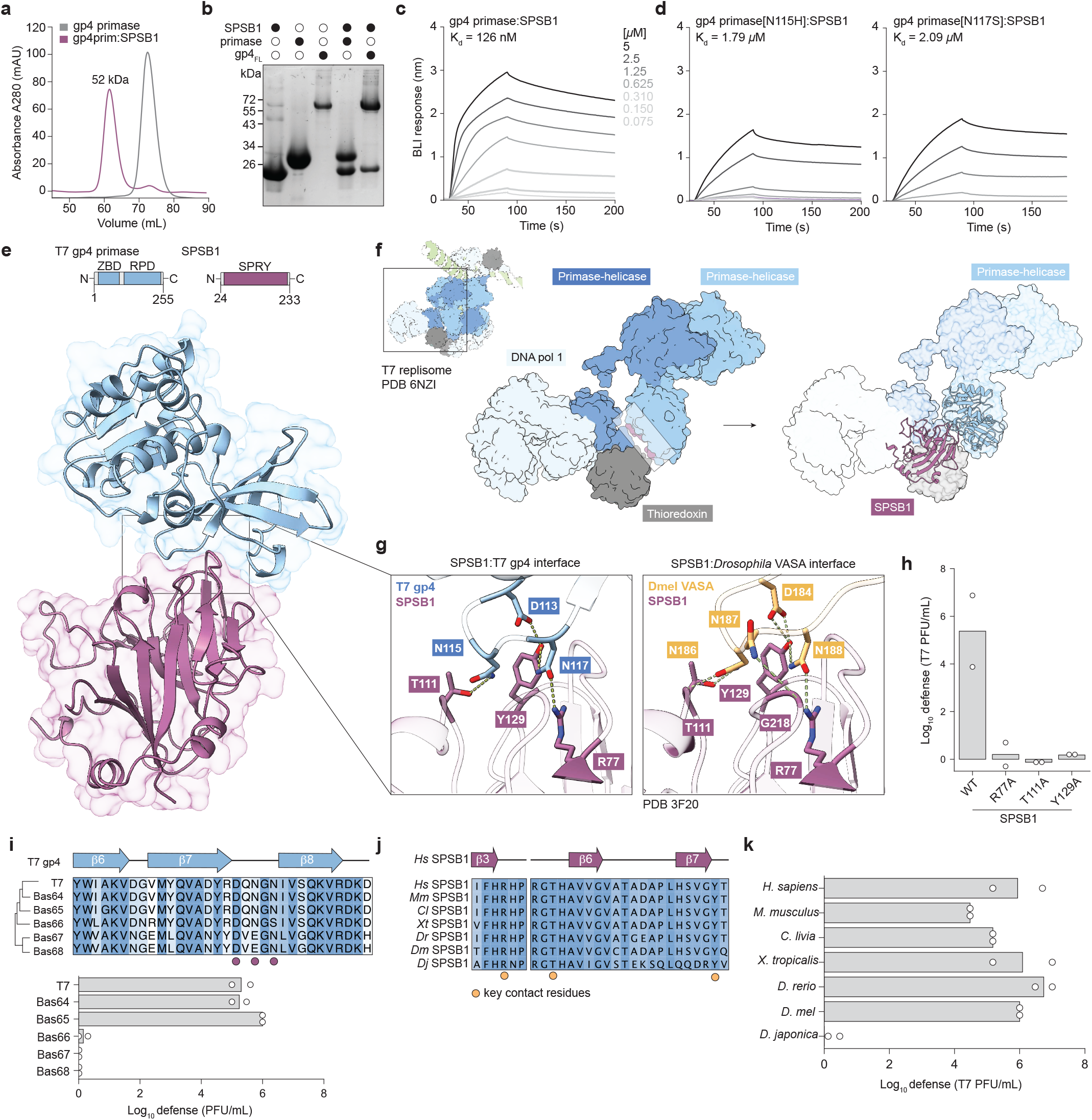
Human SPSB1 recognizes a structural DxNxN loop motif in phage T7 primase. **a**, Size-exclusion chromatography (SEC) analysis of recombinant phage T7 primase and human SPSB1–phage T7 gp4 complex. **b**, Coomassie-stained SDS-PAGE analysis of purified SPSB1, T7 gp4 primase, T7 gp4 primase-helicase (gp4FL) and SEC fractions corresponding to SPSB1:gp4 co-complexes. Open circles indicate absence while closed circles indicate presence of each protein. **c,d**, Biolayer interferometry (BLI) binding assays of immobilized 6×His-SUMO2-tagged WT gp4 primase, primase[N115H] and primase[N117S] titrated with untagged SPSB1-SPRY. Dissociation constants calculated from a 1:1 binding model using a global fit to all curves for each binding pair using vendor supplied software. **e**, Co-crystal structure of human SPSB1 in complex with phage T7 gp4 primase. Schematics indicate protein domains expressed and crystallized. **f**, SPSB1–gp4 complex overlaid onto a cryo-EM structure of the phage T7 replisome PDB 6NZI^54^. Inset shows full T7 replisome cryo-EM structure. **g**, Close-up views of SPSB1 binding pocket and conserved residues that determine recognition of phage gp4 (left) or D. mel VASA, PDB 3F20^55^ (right). Dashed green lines indicate hydrogen bonds between side-chain groups. **h**, Quantification of EOP assays in *E. coli* expressing the indicated SPSB1 variants against T7, n=2. **i**, Sequence alignment centered on gp4 loop region across phage orthologs (top). Color indicates percent consensus above 30%. T7 gp4 secondary structure is depicted above the alignment with β-sheets shown in blue. Purple circles indicate core SPSB1 motif residues. Quantification of EOP assays in SPSB1 expressing cells against Autographiviridae phages, n=2 (bottom). **j**, Sequence alignment of key SPRY domain contact residues across SPSB1 homologs from diverse animals. Color indicates percent consensus above 30%. Hs SPSB1 secondary structure is shown above with β-sheets shown in purple. Yellow circles mark critical residues involved in ligand recognition. Species names are designated as follows: Human (Hs), mouse (Mm), *Columba livia* (Cl), *Xenopus tropicalis* (Xtr), *Dan rerio* (Dr), *Drosophila melanogaster* (Dm), *Dugesia japonica* (Dj). **k**, Quantification of EOP assays in *E. coli* expressing the indicated SPSB1 homologs against T7, n=2.

To explain how a human ISG can recognize a prokaryotic viral protein, we next determined a 2.0 Å crystal structure of the SPSB1–gp4 complex (**Figure 3e** and Extended Data Table 1). In agreement with prior crystal structures^53^, the human SPSB1 SPRY domain comprises two short N-terminal helices α1 and α2 adjacent to a curved β-sandwich core formed by strands β1–β14. Five variable loops extend from the central SPSB1 β-sandwich to create a binding pocket that engages a short unstructured loop in phage T7 gp4 and positions SPSB1 on the opposite face away from the gp4 primase catalytic site. We reconstituted gp4 primase activity *in vitro* and confirmed that SPSB1 binding does not have a significant impact on RNA primer synthesis (**Extended Data Figure 4c**). Instead, comparison of the SPSB1–gp4 complex to previous structures of the phage T7 hexameric replisome assembly^54^ revealed extensive clashes between SPSB1 and the neighboring primase and DNA polymerase subunits that explain how binding would cause disruption of essential interactions required for phage T7 replication (**Figure 3f**).

The SPSB1–gp4 structure reveals a hydrophilic binding pocket in SPSB1 that reads out the sequence and spatial geometry of a loop region in gp4 located between strands β7 and β8 of the primase domain. SPSB1 contacts this loop sequence (DQNGNI) primarily at the DxNxN positions within this region. Human SPSB1 residues R77, T111, and Y129 in the binding pocket form salt-bridge interactions with phage T7 residues D113, N115, and N117 in the primase loop (**Figure 3g**), and additional main chain interactions between SPSB1 residues V216 and G218 contact the viral loop backbone and further stabilize target recognition. We introduced point mutations to SPSB1 residues R77, T111, and Y129 and confirmed that interactions in the binding pocket are necessary for phage restriction (**Figure 3h and Extended Data Figure 5a**). All phage T7 escape mutants map to the conserved viral DxNxN protein motif residues D113, N115, and N117 (**Figure 2i and 3i**). We additionally observed that Autographiviridae phages Bas66, Bas67, and Bas68 exhibit natural variation in this loop and are resistant to SPSB1 inhibition, demonstrating the importance of these positions to SPSB1 target recognition (**Figure 3i and Extended Data Figure 5b**). Previous biochemical and structural analysis of an SPSB1 homolog in *Drosophila* identified a cognate binding partner protein named VASA (and hPar4 in human cells)^53,55^. Comparison of the human SPSB1–gp4 structure with the structure of SPSB1 in complex with a peptide from *Drosophila* VASA revealed conservation of both the SPSB1 binding pocket and a shared mechanism of Asn-rich target protein loop recognition (**Figure 3g and Extended Data Figure 5c**). Sequence alignments confirmed broad conservation of the SPRY binding pocket residues in animal evolution, and we observed that diverse animal SPSB1 homologs were functional and retained the ability to inhibit phage T7 replication in bacteria (**Figure 3j,k and Extended Data Figure 5d,e**). Together, these results explain how a human ISG can protect bacteria from phage infection and highlight the conserved structural basis of SPSB1 viral protein target recognition.

### SPSB1 targets animal viral proteins for degradation

We reasoned that DxNxN loop motif recognition may underly the function of SPSB1 in the direct inhibition of animal viruses. To identify potential animal viral targets of SPSB1, we used the SPSB1–gp4 structure as a template to search a database of ∼17,000 predicted eukaryotic viral protein structures for DxNxN sequence motifs that matched the local geometry of the SPSB1 binding pocket^56,57^ (**Figure 4a and Extended Data Figure 6a**). We analyzed the top ∼100 high-confidence predictions from the *in silico* screen and identified potential enzymatic and structural viral protein targets from across 20 distinct eukaryotic viral families (**Figure 4b and Extended Data Figure 6a–d**). We co-expressed candidate proteins with SPSB1 and observed direct protein complex formation *in vitro* across diverse animal viral targets including rotavirus NTPase, NSP2^58^, human papilloma virus (HPV) DNA helicase, E1^59^, and human norovirus (HuNoV) RNA helicase, NS3^60,61^ (**Figure 4c and Extended Data Figure 6d,e**) Using size-exclusion chromatography, we confirmed that SPSB1 forms direct 1:1 complexes with animal viral targets analogous to the high-affinity interaction with phage T7 gp4 (**Extended Data Figure 6f**). Sequence alignments of homologs of eukaryotic viral protein targets revealed overall conservation of the core DxNxN protein loop motif with variation in motif residues in some viruses (such as HuNoV) potentially indicating escape from SPSB1 recognition (**Figure 4d and Extended Data Figure 6g**). Mutagenesis of DxNxN motifs in susceptible HPV E1 and HuNoV NS3 protein variants and substitution of DxNxN motifs into resistant viral variants confirmed that protein loop motif recognition is both necessary and sufficient for SPSB1 target interaction (**Figure 4e,f**). Together these results demonstrate that SPSB1 binds animal viral proteins through recognition of a common, minimal protein structural motif.

**Figure 4:**
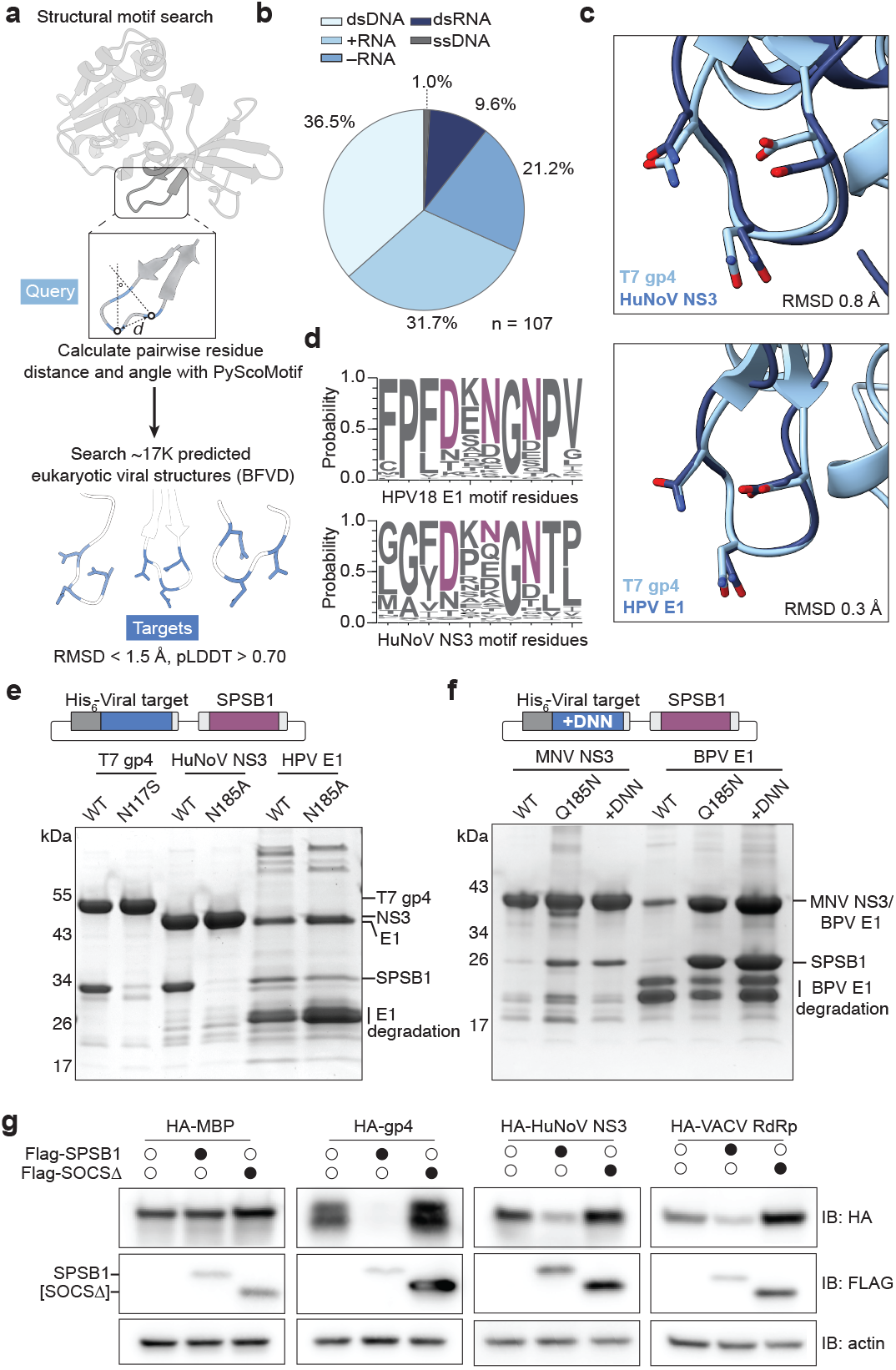
SPSB1 degrades animal viral proteins containing DxNxN motifs. **a**, Schematic of 3D structure and sequence motif search using PyScoMotif^57^. **b**, Viral species Baltimore class composition of PyScoMotif search results using gp4 loop motif as the query, n=107. **c**, Close-up views of example animal viral protein hits from PyScoMotif overlaid with T7 gp4 loop query. Human norovirus (HuNoV) NS3 (top). Human papillomavirus 18 (HPV) E1 (bottom). **d**, Sequence logos of 10 amino acid window around HPV18 E1 and HuNoV NS3 regions predicted to interact with SPSB1. Logos represent 250 homologous sequences. **e,f**, SDS-PAGE analysis of co-purified viral target variants and SPSB1-SPRY. NS3 from Murine Norovirus (MNV) and E1 from Bovine Papilloma Virus (BPV) were used to test sufficiency. +DNN mutants indicate introduction of complete DxNxN motif into the homologous region of those proteins. Schematic depicts co-expression plasmid design. **g**, Immunoblot analysis of the levels of indicated HA-tagged viral proteins co-transfected with either WT FLAG-SPSB1 or SOCS-box deficient FLAG-SPSB1 in HEK293T cells. Open circles indicate absence while closed circles indicate presence of FLAG-tagged SPSB1 variant. Actin was used as the loading control.

SPSB1 has previously been shown to regulate protein levels of cellular host targets through ubiquitin-dependent degradation^46,47,48,49^. To determine if SPSB1 can induce degradation of foreign viral proteins we constitutively co-expressed each target with SPSB1 in human HEK293T cells and measured the impact on viral target protein stability. SPSB1 expression resulted in near total degradation of phage T7 gp4 protein in cells but had no impact on a negative control protein, *E. coli* maltose-binding protein (MBP), lacking the DxNxN protein loop motif (**Figure 4g and Extended Data Figure 7a**). Testing individual candidate animal viral protein targets, we observed that SPSB1 similarly directed degradation of multiple viral targets including HuNoV NS3, a vaccinia virus poly(A) polymerase protein VP55, and a bat betacoronavirus RNA-dependent RNA polymerase protein NSP12 (**Figure 4g and Extended Data Figure 7b**). Although SPSB1 interacted with the HPV E1 helicase domain fragment^62^ *in vitro*, full length HPV E1 protein stability was unperturbed in HEK293T cells potentially indicating that SPSB1 requires cytosolic localization of viral protein targets (**Extended Data Figure 7b**). In each case, SPSB1-dependent target degradation required the SPSB1 SOCS-box domain and an intact viral DxNxN protein loop motif verifying that complex formation is required for target degradation (**Figure 4g and Extended Data Figure 7c,d**). Collectively, these data suggest a direct antiviral role for SPSB1 in degrading viral proteins containing a common sequence-specific protein loop motif.

## Discussion

Our results reveal that ISGs protect human cells by targeting common features of viral replication shared across distinct domains of life. Building on pioneering studies developing high-throughput over-expression and loss-of-function ISG screens in human cells^7,8,31,32,33,34,63,64,12^, our screen of over 300 ISGs against 11 phages in bacteria establishes a cross-kingdom viral infection model that creates unique opportunities to uncover fundamental aspects of human immunity. We show that human ISGs function as specific immune effectors in bacteria and can defend cells with potency equal to native bacterial anti-phage defense systems. Analysis of ISG activity in bacteria further demonstrates that individual human immune proteins exhibit a remarkable ability to function autonomously as antiviral restriction factors. Cross-kingdom screening also enables mechanistic insights into the function of human immune proteins by utilizing naïve viral species not previously adapted to eukaryotic antiviral defense. We show that phages can evolve to overcome human ISG-mediated immunity, driving rapid discovery of the molecular determinants of individual ISG activities. Leveraging bacteria-phage genetics, we identify human SPSB1 as a viral protein sensor that inhibits replication through steric occlusion in phages and targeted protein degradation in animal viruses. The structure of human SPSB1 in complex with a phage target protein explains how coordination of a minimal DxNxN loop motif enables recognition of a molecular feature common in both animal and bacterial viruses. SPSB1 is part of a family of four paralogs in humans, suggesting additional mechanisms of viral protein recognition likely control cellular antiviral defense. Comparative analysis of animal and bacterial immunity has provided key evolutionary insights into both the function and origin of human immune responses. Our study now extends the connection between animal and bacterial immunity beyond evolutionary comparisons and demonstrates that the powerful experimental tools of bacteria-phage models of infection can be directly applied to the study of immune protein function from any organism across the tree of life.

## Acknowledgements

The authors are grateful to members of the Kranzusch lab for helpful comments and invaluable discussion. We thank S. Hobbs, D. Wassarman, and D. Richmond-Buccola for help with phage experiments and S. Mukhopadhyay for help with cell culture experiments. Computational analyses were performed on the O2 High-Performance Compute Cluster at Harvard Medical School. The work was funded by grants to P.J.K. from the Pew Biomedical Scholars program, the Burroughs Wellcome Fund PATH program, The G. Harold and Leila Y. Mathers Charitable Foundation, the Cancer Research Institute, the Parker Institute for Cancer Immunotherapy, the Massachusetts Consortium on Pathogen Readiness (MassCPR), the Gates Foundation (INV-083469), and the National Institutes of Health (1DP2GM146250-01). S.G.F. is supported through a Cancer Research Institute Irvington Postdoctoral Fellowship (CRI14458), J.M.J.T. is supported by a Servier PhD Fellowship, S.Y. is supported by a JSPS Overseas Research Fellowships (202360072) and a Human Frontiers Science Program Long-Term Fellowship (LT0051), and A.A.R. is supported by the National Science Foundation Graduate Research Fellowship Program under Grant No. DGE 2140743. X-ray data were collected through support by an agreement between the Advanced Photon Source, a US Department of Energy (DOE) Office of Science user facility operated for the DOE Office of Science by Argonne National Laboratory under contract no. DE-AC02-06CH11357, through the Northeastern Collaborative Access Team beamlines, which are funded by the National Institute of General Medical Sciences from the National Institutes of Health (P30 GM124165) and a NIH-ORIP HEI grant (S10OD021527).

## Author contributions

The study was designed and conceived by S.G.F. and P.J.K. All phage challenge, genetic experiments and bioinformatics were performed by S.G.F. Primase activity assays were performed by J.M.J.T. Biochemical and structural biology experiments were performed by S.G.F. with assistance from J.E.H. and S.Y. BLI protein interaction analysis was performed by S.G.F. with assistance from A.A.R. and A.G.S. The manuscript was written by S.G.F and P.J.K. All authors contributed to editing the manuscript and support the conclusions.

## Additional Information

Correspondence and requests for materials should be addressed to P.J.K. All illustrations were created using Adobe Illustrator.

## Data Availability Statement

Coordinates and structure factors of the Human SPSB1-SPRY domain in complex with phage T7 gp4 primase have been deposited in Protein Data Bank (PDB) under the accession code 36FM. All other data are available in the manuscript or the supplementary information.

## Materials and Methods

### Bacterial strain and growth conditions

*E. coli* TOP10 (Invitrogen) were used for all plasmid cloning and were grown at 37°C in Luria-Bertani (LB) media supplemented with appropriate antibiotic (100 µg/mL ampicillin, 34 µg/mL chloramphenicol). *E. coli* BW25113 was used for propagating phage stocks and in all phage challenge assays. E. coli BL21-RIL or BL21-LBSTR cells were used for protein expression (Agilent).

### Efficiency of plating assays

Ten dsDNA *E. coli* phages from the BASEL collection^66^ and one ssDNA phage (SECφ17) were selected for phage challenge assays. Phage stocks were plaque purified twice and phage identity was confirmed either through PCR-based genotyping or by whole-genome sequencing. Full length human ISG ORFs were synthesized and cloned into an arabinose-inducible pBAD vector (GenScript). *E. coli* BW25113 cells were transformed with human ISG plasmids and plated onto LB supplemented with 1% glucose and 100 µg/mL ampicillin. Single colonies were inoculated into LB containing 100 µg/mL ampicillin and grown for 3-4 h at 37°C to OD600 = 0.8–1.0. Cultures were then diluted to OD600 = 0.2-0.3 into 1% LB-low melt agar containing a final concentration of 5mM MgCl_2_, 5mM CaCl_2_, 0.1 mM MnCl_2_, 0.2 % arabinose and 100 µg/mL ampicillin. Top agar containing cultures were then plated onto solid LB media containing 0.2% arabinose and 100 µg/mL ampicillin and allowed to set for 30 min. 10-fold serial dilutions of phage stocks were prepared in SM Buffer (100 mM NaCl, 50 mM Tris-HCl pH 7.5, and 8 mM MgSO_4_) and 2.5 µL of each dilution was spotted onto each plate. Droplets were allowed to dry for 20 min and incubated at 30°C. Fold changes were calculated as the ratio between plaque-forming units per milliliter (PFU/mL) of phages plated on GFP-expressing bacteria and human ISG-expressing bacteria. Plaques were imaged on a ChemiDoc MP Imaging System.

### Bacterial toxicity assays

Human ISG plasmids were transformed into E. coli BW25113 and plated onto LB agar supplemented with 1% glucose and 100 µg/mL ampicillin to repress expression from the arabinose-inducible promoter. One colony from each transformation was picked into 4 mL LB with 1% glucose and 100 µg/mL ampicillin and grown overnight at 37°C, shaking at 230 RPM. Cells were centrifuged at 3,220 × g for 5 min at RT then resuspended in 3 mL PBS. 10-fold serial dilutions of cultures were prepared in PBS and incubated for 15 min at 37°C. 5 µL of each dilution was spotted onto LB plates supplemented with either 1% glucose and 100 µg/mL ampicillin or 0.2% arabinose and 100 µg/mL ampicillin. Plates were then incubated at 37°C overnight and imaged on a ChemiDoc MP Imaging System.

### Recombinant protein expression and purification

All proteins were cloned into custom pET vectors as N-terminal 6×His-SUMO2 fusion proteins with a GS linker and transformed into *E. coli* strain BL21-LBSTR^67^. Colonies for protein expression were grown in starter MDG media cultures (0.5% glucose, 25 mM Na2HPO4, 25 mM KH2PO4, 50 mM NH4Cl, 5 mM Na2SO4, 2 mM MgSO4, 0.25% aspartic acid, 100 µg/mL ampicillin, 34 µg/mL chloramphenicol and trace metals)^68^ and grown overnight at 37°C. MDG cultures were used to seed 1L M9ZB (0.5% glycerol, 1% Cas-Amino Acids, 47.8 mM Na2HPO4, 22 mM KH2PO4, 18.7 mM NH4Cl, 85.6 mM NaCl, 2 mM MgSO4, 100 µg/mL ampicillin, 34 µg/mL chloramphenicol and trace metals) cultures and grown at 37°C until OD600 1.5–2.5. Recombinant protein expression was then induced with 0.5 mM IPTG and incubated at 16°C with shaking for ∼17 h before harvesting. Bacterial pellets were resuspended and sonicated in cold lysis buffer (20 mM HEPES-KOH pH 7.5, 400 mM NaCl, 30 mM imidazole and 10% glycerol, 1 mM DTT) and purified using Ni-NTA resin (Qiagen). Ni-NTA resin was washed with lysis buffer with 1 M NaCl and eluted with lysis buffer supplemented with 300 mM imidazole. Elution samples were treated with ∼200 µg recombinant hSENP2 protease to remove the SUMO2 tag and dialyzed overnight at 4°C into 20 mM HEPES-KOH pH 7.5, 250 mM KCl, 1 mM DTT in 14 kDa molecular weight cutoff dialysis tubing (Ward’s Science). Dialyzed samples were then concentrated using a 10 kDA MWCO centrifugal filter (Millipore Sigma) then separated by size-exclusion chromatography in 20 mM HEPES-KOH pH 7.5, 250 mM KCl, 1 mM TCEP gel filtration buffer using a 16/600 Superdex 75 column (Cytiva).

### Co-purification and His pull-down assays

Recombinant human SPSB1 (NP079382.2) and each viral protein target were cloned into custom pET co-expression vectors where, unless noted otherwise, SPSB1 was untagged and each viral protein was expressed as a N-terminal 6×His-SUMO2 tag fusion^67^. Co-expression plasmids were expressed as described above. 6×His-SUMO2 tagged viral proteins were purified using Ni-NTA resin as described above except that bound protein complexes were washed in lysis buffer with 400 mM NaCl and eluted into 20 mM HEPES-KOH pH 7.5, 200 mM NaCl, 1 mM DTT.

### Phage escape isolation and mutant sequencing analysis

To isolate phage escape mutants, EOP assays were performed as described as above using the indicated ISGs expressed in *E. coli* BW25113. Single plaques were isolated from ISG-expressing cells into 50 µL SM buffer. Single plaques were propagated and re-purified on naïve BW25113 cells. Genomic DNA extraction and whole genome sequencing of parental and escape phage stocks was carried out by SeqCenter (Pittsburgh PA, USA) using paired-end Illumina sequencing. Paired-end reads were aligned using Bowtie 2.2.4^69^. Variants were identified using Bcftools 1.21^70^ assuming a ploidy of 1 and default settings. Mutations were filtered for positions where there was at least 150× read depth, a Phred base quality score of at least 25, and were not present in the parental genome.

### Biolayer interferometry

Biolayer interferometry (BLI) experiments were performed using a BLItz instrument (Fortebio) with Ni-NTA biosensors (Fortebio). Viral proteins were immobilized to Ni-NTA biosensors and SPSB1-SPRY was used as the analyte. Proteins immobilized for BLI were expressed as N-terminal 6×His-SUMO2 tag fusions and purified as described above with addition of 10% glycerol to the gel filtration buffer. Analyte samples were purified as described above and cleaved with hSENP2 to remove 6×His-SUMO2. All samples were diluted in 20 mM HEPES-KOH pH 7.5, 10% glycerol, 100 mM KCl, 1 mM TCEP-KOH. Titrations starting at 5 µM and at least 5 additional concentrations were performed and dissociation constants (K_d_) were determined by applying a global fit assuming a 1:1 binding isotherm using vendor-supplied software.

### Crystallization and structure determination

Crystals of human SPSB1 residues 24–233 and phage T7 gp4 primase residues 10–255 were grown in a hanging-drop format using the hanging-drop vapor diffusion method for 3–7 days at 18°C. Recombinant human SPSB1 and T7 primase were mixed in a 1:1 molar ratio and diluted to 9 mg/mL in a buffer containing 20 mM HEPES-KOH pH 7.5 and 125 mM KCl. The protein mixture was allowed to equilibrate to 18°C for 15 min and screened in 96-well trays containing 70 µL of reservoir solution (NeXtal) and 0.2 µL drops. Resultant crystals were further optimized by mixing purified complex and reservoir solution (0.1 M HEPES pH 7.4, 0.1 M NaCl, 1.3 M (NH4)_2_SO_4_) in a 1:1 ratio. Crystals were cryo-protected with a reservoir solution supplemented with 10% ethylene glycol and harvested by flash freezing in liquid nitrogen.

X-ray diffraction data were collected at the Advanced Photon Source Argonne National Laboratory (APS). Data were collected using the NECAT-remote GUI v6.2.3 and processed with RAPD2^71^ and AIMLESS 1.12.16^72^. Experimental phase information was determined by molecular replacement using Phaser-MR in Phenix^73^ with prior crystal structures of the phage T7 primase (PDB 1NUI) and human SPSB1 (PDB 2JK9). Model building was carried out in Coot 0.9.8.96^74^ and refinement was done using Phenix 2.1^73^. A summary of crystallographic statistics is provided in Extended Data Table 1. All structural figures were generated using ChimeraX^75^.

### Primase activity assay

To measure RNA synthesis, 2 µM of WT primase and full length mutant [A247T] gp4 and 20 µM of SPSB1 were incubated with 4 µM of ssDNA template^50,76^: 5′–CAGTGACGGGTCGTTTATCGTCGGCA–3′ or 5′– T_20_ATTCGTAATCTGCAGGCATGGTCAAT_5_ATAGAGTTAT–3′ on ice for 60 mins. Protein–DNA mixture was incubated with unlabeled 1000 µM ATP (Jena Bioscience) and 0.5 µM α^32^P-labeled CTP (Perkin Elmer / Revvity, approximately 1 µCi) in reaction buffer (50 mM Tris-HCl pH 7.5, 10 mM MgCl_2_, 10 mM DTT, 100 µg/ml BSA) at 37°C for 60 mins. Reactions were terminated with addition of 0.5 µl of Quick CIP phosphatase (New England Biolabs) at 37°C for 60 mins to remove terminal phosphate groups from unreacted nucleotides. Where indicated, reactions were additionally treated with 0.5 µl nuclease P1 (NEB) or 0.5 µl RNase T1 (ThermoFisher) and 0.5 µl RNase A (NEB). Each reaction was denatured with 2× STOP loading dye (95% (v/v) deionized formamide, 20 mM EDTA, 0.01% (v/v) bromophenol blue and xylene cyanol) and analyzed on a 20% Urea-PAGE gel and exposed to a phosphor-screen before imaging with a Typhoon Trio Variable Mode Imager (GE Healthcare).

### Sequence and structure motif search

Animal viral protein models were obtained from the Big Fantastic Viral Database (BFVD)^56^ and filtered for viruses with human hosts. *In silico* sequence structure motif searches were done using PyScoMotif^57^. 17,633 animal viral proteins were indexed using PyScoMotif and search parameters were set such that target motifs contained only DxNxN residues in any order.

### Cell culture and transient transfection assays

Human HEK293T cells were grown in DMEM supplemented with GlutaMAX (Thermo Fisher Scientific) and 10% FBS. All cell lines were obtained from the ATCC and grown at 37°C and 5% CO_2_. Full length ORFs corresponding to Human SPSB1 (NP079382.2), HuNoV NS3 (QBF80211.1), VACV VP55 (AAB59821.1), HPV E1 (WWB08199.1), bat βCoV RdRP (KF636756.1), RV3 RdRP (AWD85050.1), phage T7 gp4 (NP041977.1) were cloned into constitutive pCMV vectors. Transient cotransfections were carried out using X-tremeGene HP DNA Transfection Reagent (Roche) using 1 µg of DNA per plasmid. Cells for Western blot experiments were harvested at 48 h post-transfection.

### Western Blotting

Transiently transfected cells were collected by centrifugation (500 × g for 5 min at 4°C) washed once with ice-cold PBS, centrifuged again and lysed with 1% Triton X-100 buffer (10 mM HEPES-KOH pH 7.5, 10 mM KCl, 3 mM MgCl_2_, 10% glycerol, 1 mM DTT, 1% Triton X-100), nutating at 4°C for 30 min. Lysates were clarified by centrifugation (20,000 × g for 15 min at 4°C). Protein lysates were separated on 4–20% TGX gels (Bio Rad) in Tris-Glycine buffer then transferred onto PVDF membranes. Membranes were blocked with 5% milk in TBST (0.05% Tween-20) for 1 h at RT. Primary antibodies were incubated overnight at 4°C and secondary antibodies for 1 h at RT. FLAG epitope tags were probed by a primary FLAG antibody (1:2000, #14793; Cell Signaling Technology), HA epitope tags were probed by a primary HA antibody (1:3000, #3724; Cell Signaling Technology) and β-actin loading controls were probed by a primary β-actin conjugated to HRP (1:4000, #4967; Cell Signaling Technology). An HRP-conjugated anti-rabbit IgG (1:2000-1:4000, #7074; Cell Signaling Technology) was used as a secondary antibody against all primary antibodies. All blots were developed with either SuperSignal West Dura Extended Duration Substrate (Thermo Fisher Scientific) or ECL Select (Cytiva) and visualized on a ChemiDoc MP Imaging System.

### Statistics and reproducibility

Experimental details regarding replicates and sample size are described in the figure legends.

---

**Extended Data Figure 1:**
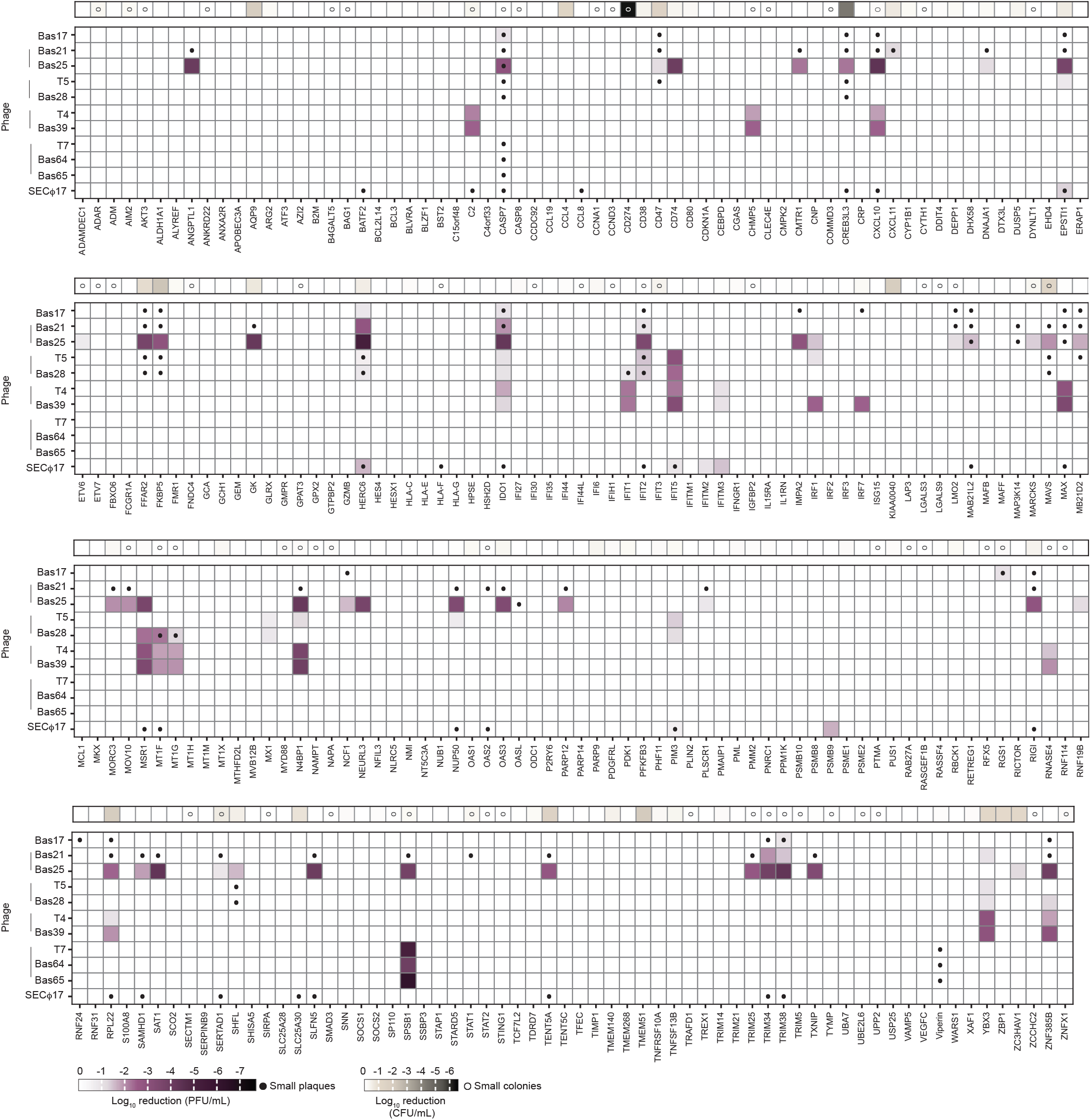
Systematic discovery of anti-phage activity in 255 human ISGs. **a**, Heatmap of reduction in phage replication (PFU/mL) upon arabinose inducible expression of 255 human ISGs against 11 bacteriophages. 51 ISGs were removed from further characterization due to toxicity. Viral titer measurements are log_10_ transformed and replication foldchanges reflect measurements relative to titers in GFP-expressing control cells. Closed circles indicate small plaque size morphology whereas lines next to phage labels indicate family membership. Heatmap of reduction in *E. coli* host cell growth in colony forming units per mL (CFU/mL) are aligned at the top of each row. Open circles indicate small colony formation phenotypes. Data represent mean of two independent replicates.

**Extended Data Figure 2:**
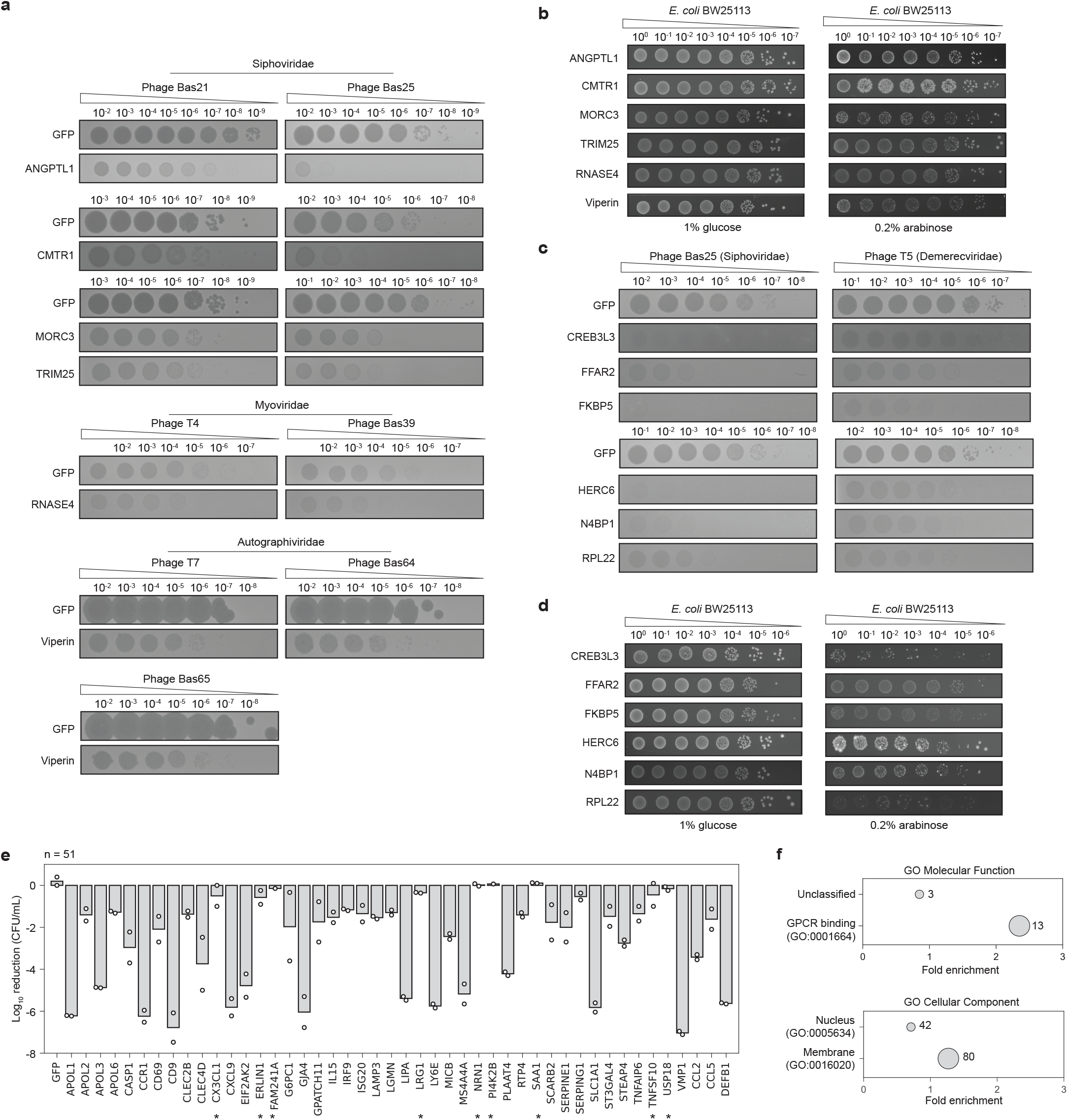
Characterization of human ISG anti-phage activity and overexpression toxicity. **a**, Representative EOP assays of serially diluted phages plated on *E. coli* expressing a GFP control or the indicated human ISGs. Phage family membership is indicated above respective phages. **b**, Representative toxicity spot assays of ISGs tested in (a). *E. coli* transformed with the indicated ISGs were serially diluted on LB supplemented with either 1% glucose (promoter off) or 0.2% arabinose (promoter on). **c**, Representative EOP assays of serially diluted phages plated on *E. coli* expressing a GFP control or the indicated toxic human ISGs. Phage family is indicated in parentheses. **d**, Representative toxicity spot assays of ISGs tested in (c). **e**, Quantification of toxicity assays of human ISGs where EOP assays were unquantifiable. Overall, 51 ISGs strongly reduced cell viability, n=2. Asterisks indicate ISGs that induced severe small colony phenotypes. **f**, Gene-ontology (GO) terms associated with ISGs that reduced CFU/mL by at least 10-fold or reduced colony size. Enriched GO terms for Molecular Function (top) or for Cellular Component (bottom). Bubble size indicates number of ISGs (Fisher’s exact test, Bonferroni’s correction, p-values < 0.05). Enriched GO annotations for membrane compartments and GPCR binding suggest that these ISGs contained transmembrane segments toxic to *E. coli*.

**Extended Data Figure 3:**
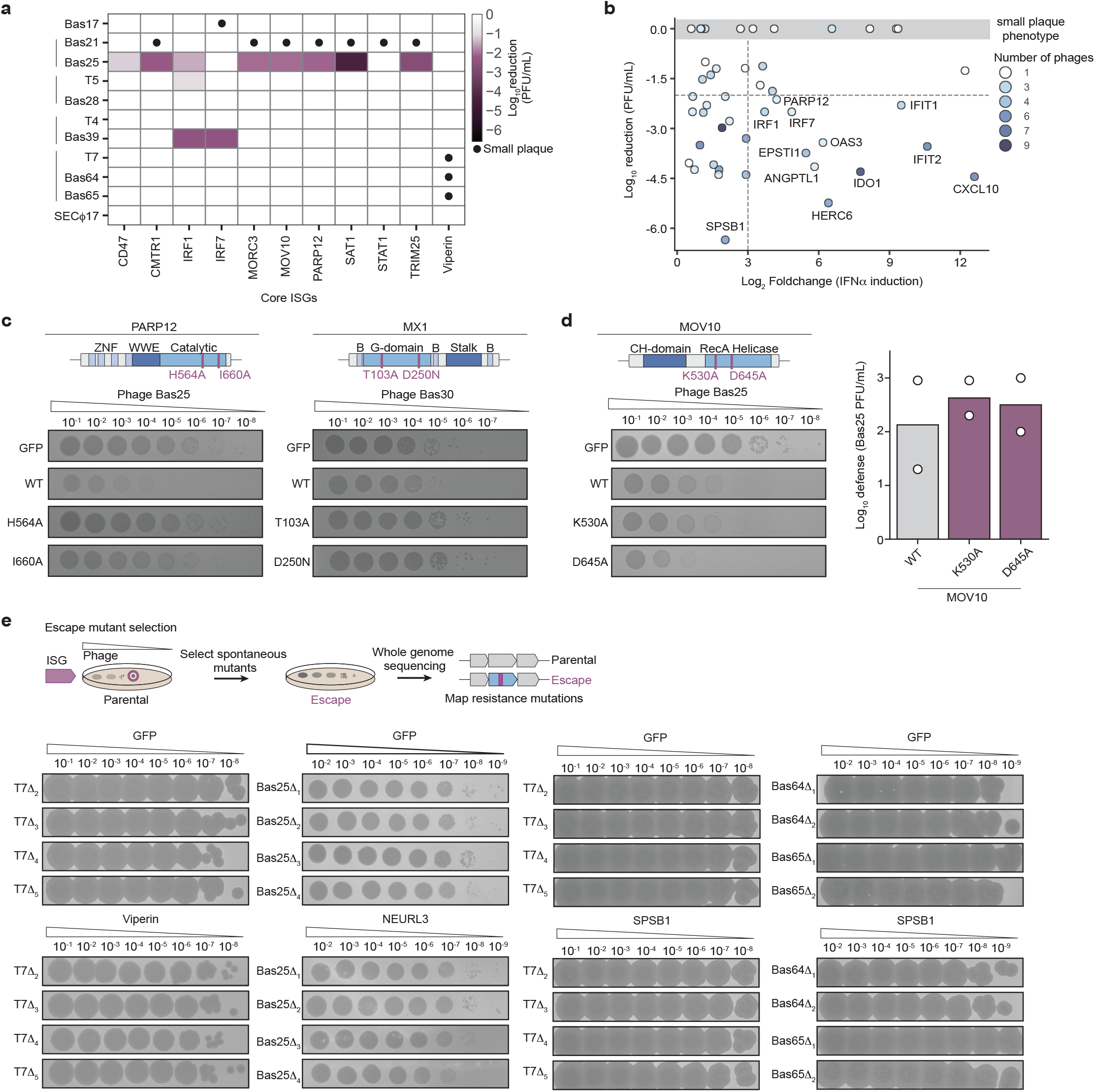
Characterization of human ISG anti-phage activity and overexpression toxicity. **a**, Heatmap of log_10_ reduction in phage titer (PFU/mL) for core ISGs in **Figure 2c**. **b**, Plot of phage defense (reduction in PFU/mL) against mRNA expression after 4h of IFNα treatment expressed as log_2_ foldchanges relative to mock-treated cells^5^. Dashed lines demarcate 100-fold reduction in phage titer and 8-fold increase in IFN-dependent mRNA expression. Colors represent number of phages susceptible to each ISG. We observed a very weak, positive correlation between ISG mRNA expression upon IFNα activation and our defense phenotypes, suggesting that IFN induction levels are not predictive of defense against phages. **c**, Representative EOP assays of *E. coli* expressing indicated ISG and mutant variants. Schematics show protein domain architecture of each ISG with positions of mutations indicated in purple. **d**, As in (c) but for the ISG, MOV10 (left). Quantification of MOV10 EOP assays against Bas25, n=2. e, Schematic of strategy for selecting spontaneous phage escape mutants (top). Representative EOP assays of *E. coli* expressing GFP or indicated ISGs challenged with purified escape phages. Numbers indicate separate clones.

**Extended Data Figure 4:**
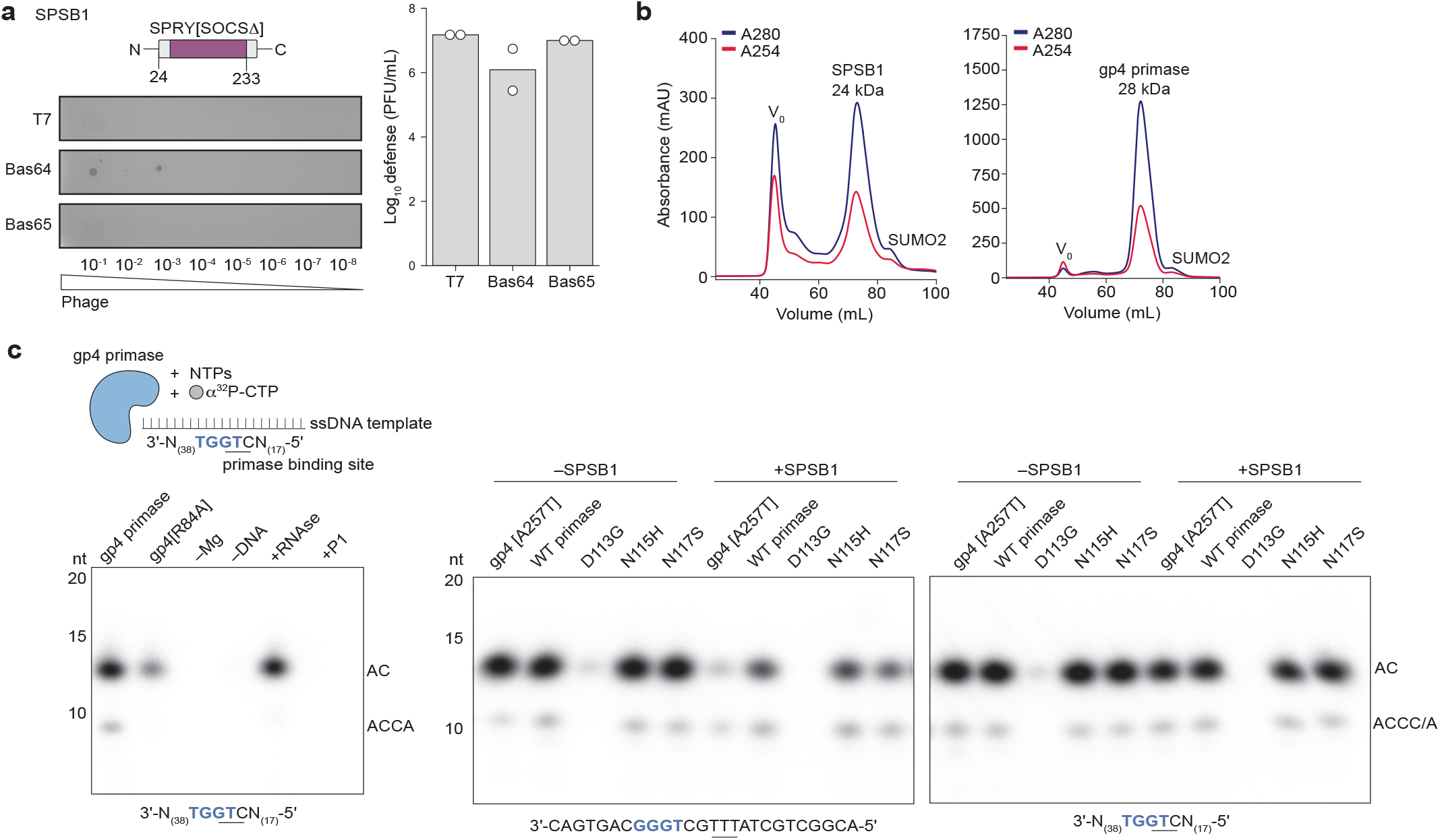
Biochemical characterization of human SPSB1 inhibition of phage T7 primase activity. **a**, Representative EOP assay of *E. coli* expressing SPSB1 with a deletion of the SOCS-box domain challenged with the indicated phages (left). Quantification of EOP assays relative to GFP control cells, n=2 (right). Schematic represents SPSB1 mutant domain architecture. **b**, Size exclusion chromatography traces of purified 6×His-SUMO2-SPSB1 and 6×His-SUMO2-gp4 primase. Void volume (V_0_) and SUMO2 cleavage products are indicated. **c**, Urea-PAGE analysis of impact of SPSB1 on phage T7 gp4 RNA primer synthesis activity using NTPs, α32P-labeled CTP and the indicated ssDNA primer template. Expected priming site is underlined. A gp4 primase catalytic mutant, R84A, was used as a negative control. Verification of RNA primer synthesis of purified gp4 primase (left). Effect of SPSB1 on gp4 primase activity (middle, right). Two ssDNA templates were tested in primase reactions—a minimal template (middle) and a long template (right). SPSB1 reduces primase activity on minimal templates. A full length gp4 primase-helicase (containing a mutant, A257T, in linker region that was used for expression) was used as a positive control for synthesis. D113G, N115H and N117S lanes correspond to recombinant primase engineered with respective escape mutations. Migration of RNA products is inversely correlated with RNA size ladder due to loss of triphosphates upon CIP-treatment to terminate reactions^65^.

**Extended Data Figure 5:**
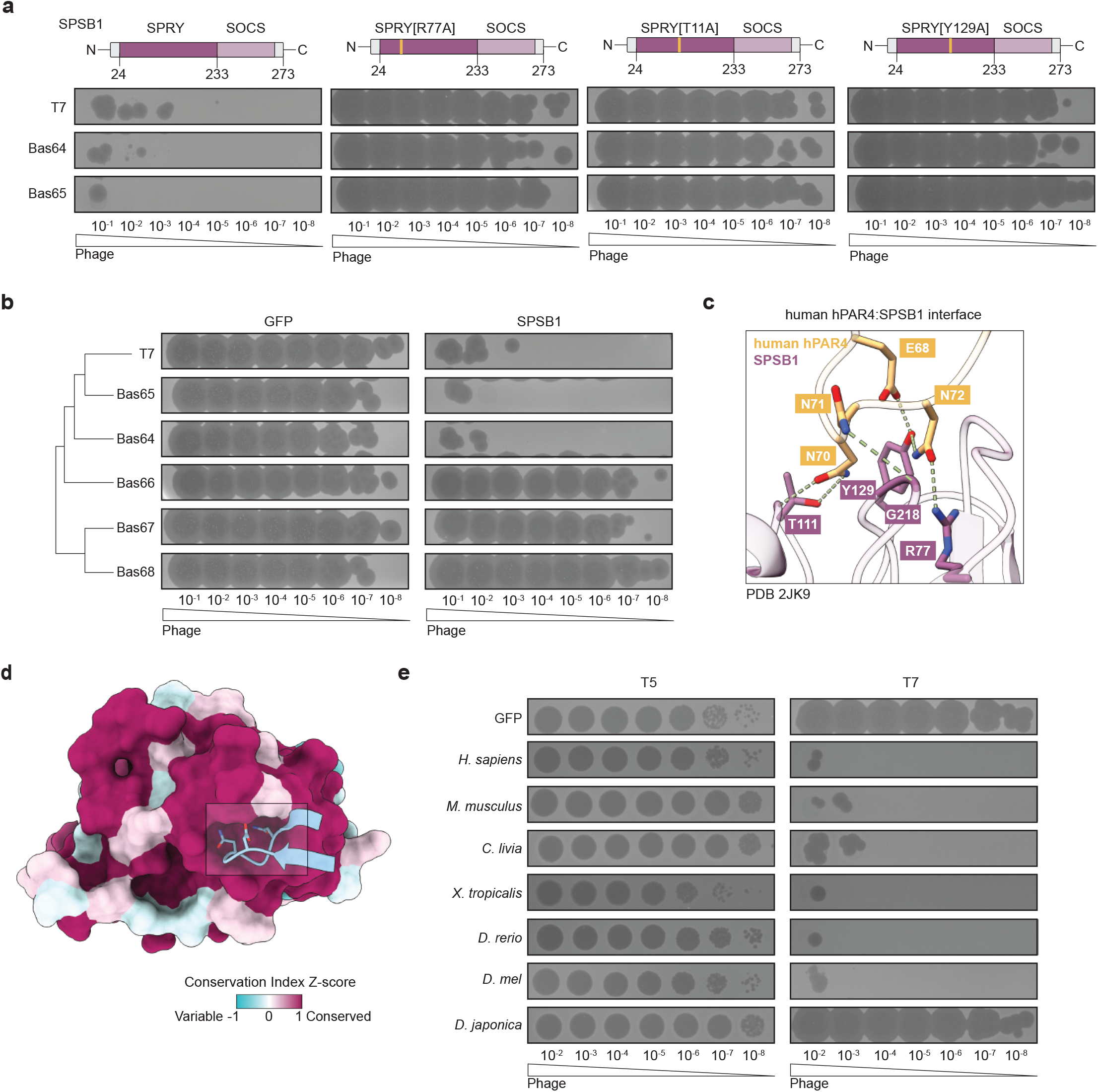
Human SPSB1 binding pocket is conserved. **a**, Representative EOP assays of *E. coli* expressing indicated SPSB1 variants challenged with phage T7, Bas64 and Bas65 as in **Figure 3h**. Schematics depict SPSB1 protein domain architecture with each mutation in the SPRY domain indicated in yellow. **b**, Representative EOP assays of SPSB1-expressing *E. coli* challenged with Autographiviridae phages as in **Figure 3i**. **c**, Close-up view of human SPSB1 in complex with human hPar4 PDB 2JK9. Dashed green lines indicate hydrogen bonds between side-chain groups. **d**, Surface representation of SPSB1-gp4 crystal structure. Surface is colored by degree of conservation calculated as the Z-score of the conservation index (AL2CO entropy score) from 250 SPSB1 homolog sequences. Phage T7 gp4 interacting peptide loop is colored in blue. **e**, Representative EOP assays of *E. coli* expressing SPSB1 animal homologs challenged with either phage T5 which is not susceptible to SPSB1 and phage T7 as in **Figure 3k**.

**Extended Data Figure 6:**
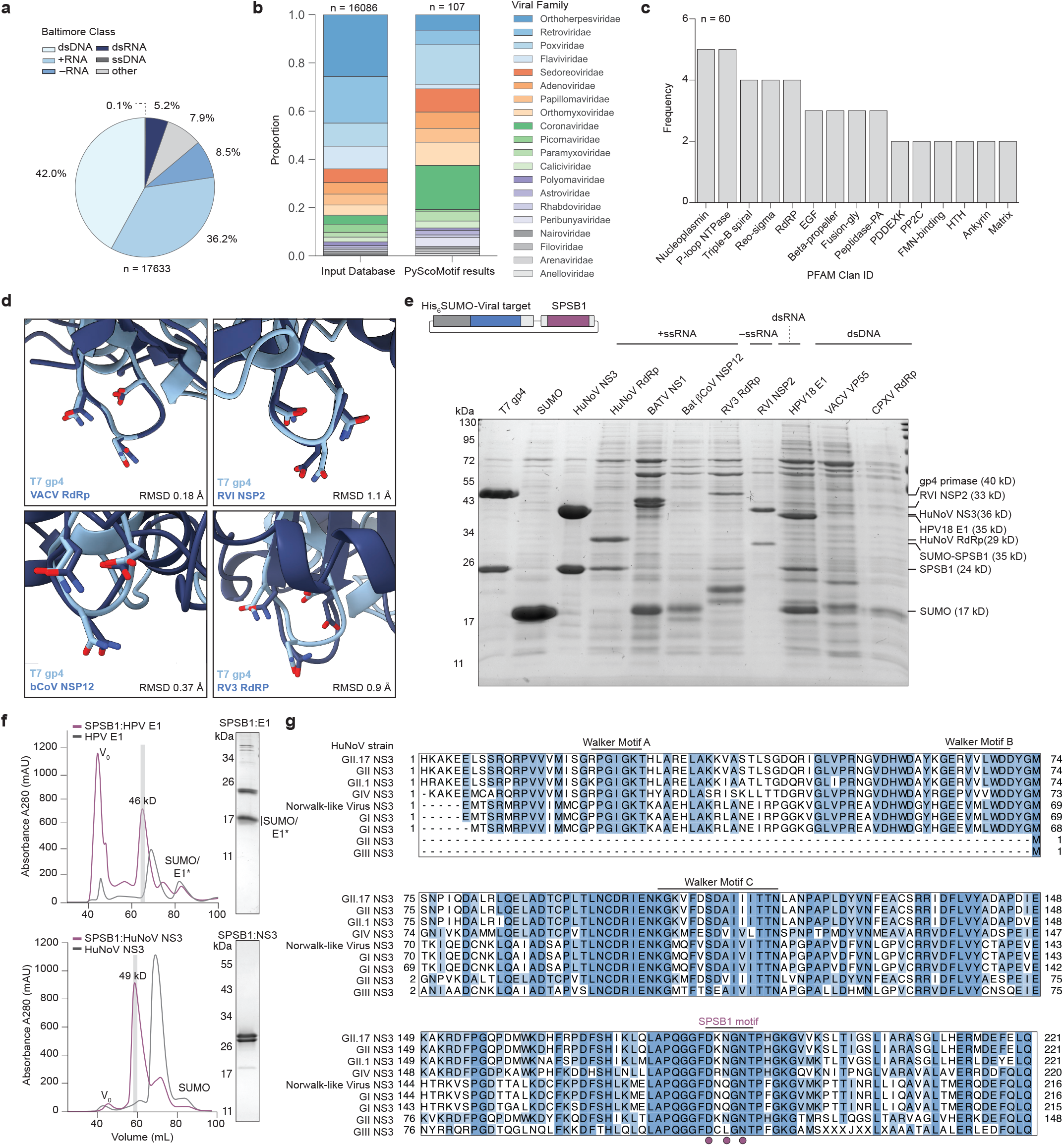
Phage T7 gp4 DxNxN motif is found in animal viral proteins. **a**, Viral species Baltimore class composition of database of AF2 predicted animal viral protein structures from BFVD, n=17633 predicted structures. **b**, Comparison of viral family composition of proteins predicted to have a DxNxN motif (PyScoMotif results) versus viral proteins used in search database (Input Database). **c**, Number of occurrences of annotated PFAM Clan IDs across PyScoMotif search hits with pLDDT scores above 0.70, n=60. Only Clan IDs present more than once are depicted. **d**, Close-up views of select PyScoMotif search hits overlaid with DxNxN motif from gp4. Hits are from vaccinia virus (VACV) and rotavirus I (RVI) (top) and bat betacoronavirus (βCoV) and respirovirus 3 (RV3) (bottom). **e**, Coomassie-stained SDS-PAGE analysis of co-purified animal viral proteins and SPSB1-SPRY. Animal viral proteins were tagged with 6×His-SUMO2 and SPSB1 was untagged except in the case of RVI-NSP2 which was untagged and 6×His-SUMO2-SPSB1 was used as bait. Schematic depicts co-expression plasmid design. Baltimore class designation is indicated above each virus. **f**, SEC profile overlay of recombinant HPV E1 expressed alone or with SPSB1 SPRY (top). SEC profile overlay of recombinant HuNoV NS3 expressed alone or with SPSB1 (bottom). SDS-PAGE analysis taken from SPSB1:E1 and SPSB1:NS3 complex fractions (right). **g**, Sequence alignment of NS3 RNA helicase sequences taken from 5 representative strains (GI, GII, GII, GIV, Norwalk-like virus). Canonical SF3 helicase motifs (Walker motifs A,B,C) are indicated. SPSB1 motif residues are indicated as purple circles.

**Extended Data Figure 7:**
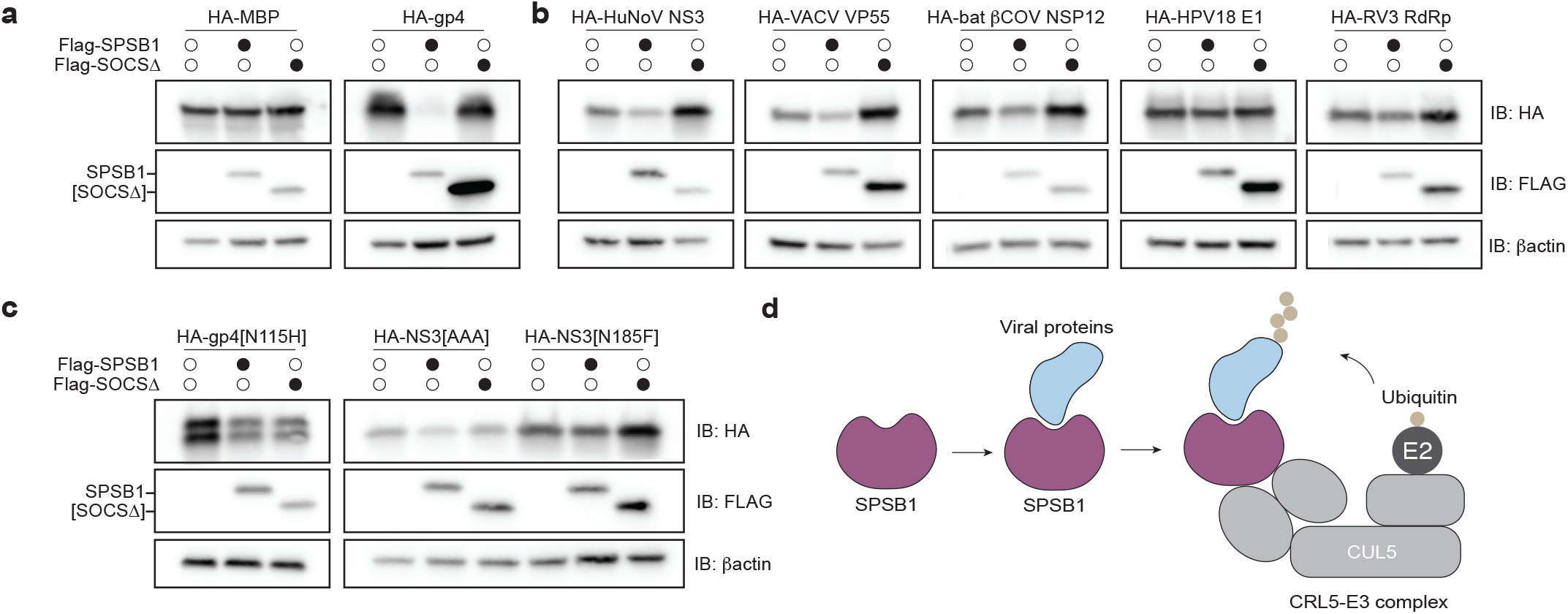
SPSB1 degrades animal virus proteins in a DxNxN motif-dependent manner. **a**, Immunoblot analysis of the levels of HA-tagged animal viral proteins or FLAG-tagged SPSB1 variants in HEK293T cells. Open circles indicate absence of co-transfected SPSB1 variant while closed circles indicate presence. Actin was used as the loading control. Western blots represent an independent replicate from what is shown in **Figure 4g**. **b**, As in (a) for the indicated HA-tagged animal viral proteins. Western blots of NS3 and VP55 represent an independent replicate from what is shown in **Figure 4g**. **c**, Immunoblot analysis as in (a,b) of HA-tagged mutants of phage T7 gp4 (left) or HuNoV NS3 (right). The NS3[AAA] mutant contains alanine substitutions in the core DxNxN residues (AxAxA). **d**, Model of SPSB1-mediated antiviral immunity.

## References

1. Knight, E., Jr & Korant, B. D. Fibroblast interferon induces synthesis of four proteins in human fibroblast cells. Proc. Natl. Acad. Sci. U. S. A. 76, 1824–1827 (1979).

2. de Veer, M. J. et al. Functional classification of interferon-stimulated genes identified using microarrays. J. Leukoc. Biol. 69, 912–920 (2001).

3. Müller, U. et al. Functional role of type I and type II interferons in antiviral defense. Science 264, 1918–1921 (1994).

4. Schneider, W. M., Chevillotte, M. D. & Rice, C. M. Interferon-stimulated genes: a complex web of host defenses. Annu. Rev. Immunol. 32, 513–545 (2014).

5. Shaw, A. E. et al. Fundamental properties of the mammalian innate immune system revealed by multispecies comparison of type I interferon responses. PLoS Biol. 15, e2004086 (2017).

6. McDougal, M. B., Boys, I. N., De La Cruz-Rivera, P. & Schoggins, J. W. Evolution of the interferon response: lessons from ISGs of diverse mammals. Curr. Opin. Virol. 53, 101202 (2022).

7. Schoggins, J. W. et al. A diverse range of gene products are effectors of the type I interferon antiviral response. Nature 472, 481–485 (2011).

8. Schoggins, J. W. et al. Pan-viral specificity of IFN-induced genes reveals new roles for cGAS in innate immunity. Nature 505, 691–695 (2014).

9. García-Sastre, A. Ten strategies of interferon evasion by viruses. Cell Host Microbe 22, 176–184 (2017).

10. Feng, H., Zhang, Y.-B., Gui, J.-F., Lemon, S. M. & Yamane, D. Interferon regulatory factor 1 (IRF1) and anti-pathogen innate immune responses. PLoS Pathog. 17, e1009220 (2021).

11. Schoggins, J. W. Interferon-Stimulated Genes: What Do They All Do? Annu Rev Virol 6, 567–584 (2019).

12. Krey, K. et al. The ISG Atlas: a loss-of-function analysis characterizes antiviral properties of interferon stimulated genes. Nat. Commun. 17, 4206 (2026).

13. Der, S. D., Zhou, A., Williams, B. R. & Silverman, R. H. Identification of genes differentially regulated by interferon alpha, beta, or gamma using oligonucleotide arrays. Proc. Natl. Acad. Sci. U. S. A. 95, 15623–15628 (1998).

14. Whiteley, A. T. et al. Bacterial cGAS-like enzymes synthesize diverse nucleotide signals. Nature 567, 194–199 (2019).

15. Cohen, D. et al. Cyclic GMP-AMP signalling protects bacteria against viral infection. Nature 574, 691–695 (2019).

16. Ye, Q. et al. HORMA domain proteins and a Trip13-like ATPase regulate bacterial cGAS-like enzymes to mediate bacteriophage immunity. Mol. Cell 77, 709–722.e7 (2020).

17. Burroughs, A. M. & Aravind, L. Identification of uncharacterized components of prokaryotic immune systems and their diverse eukaryotic reformulations. J. Bacteriol. 202, e00365–20 (2020).

18. Bernheim, A. et al. Prokaryotic viperins produce diverse antiviral molecules. Nature 589, 120–124 (2021).

19. Gao, L. A. et al. Prokaryotic innate immunity through pattern recognition of conserved viral proteins. Science 377, eabm4096 (2022).

20. Kibby, E. M. et al. Bacterial NLR-related proteins protect against phage. Cell 186, 2410–2424.e18 (2023).

21. Johnson, A. G. et al. Bacterial gasdermins reveal an ancient mechanism of cell death. Science 375, 221–225 (2022).

22. van den Berg, D. F. et al. Bacterial homologs of innate eukaryotic antiviral defenses with anti-phage activity highlight shared evolutionary roots of viral defenses. Cell Host Microbe 32, 1427–1443.e8 (2024).

23. Cury, J. et al. Conservation of antiviral systems across domains of life reveals immune genes in humans. Cell Host Microbe 32, 1594–1607.e5 (2024).

24. Wein, T. et al. CARD domains mediate anti-phage defence in bacterial gasdermin systems. Nature 639, 727–734 (2025).

25. Bonhomme, D. et al. A human homolog of SIR2 antiphage proteins mediates immunity via the Toll-like receptor pathway. Science 389, eadr8536 (2025).

26. Perez Taboada, V. et al. Bacterial Schlafen proteins mediate phage defence. Nat. Microbiol. 1–12 (2026) doi:10.1038/s41564-026-02277-8.

27. Doron, S. et al. Systematic discovery of antiphage defense systems in the microbial pangenome. Science 359, eaar4120 (2018).

28. Richmond-Buccola, D. et al. A large-scale type I CBASS antiphage screen identifies the phage prohead protease as a key determinant of immune activation and evasion. Cell Host Microbe 32, 1074–1088.e5 (2024).

29. Lowey, B. et al. CBASS immunity uses CARF-related effectors to sense 3’-5’- and 2’-5’-linked cyclic oligonucleotide signals and protect bacteria from phage infection. Cell 182, 38–49.e17 (2020).

30. Kane, M. et al. Identification of Interferon-Stimulated Genes with Antiretroviral Activity. Cell Host Microbe 20, 392–405 (2016).

31. Feng, J. et al. Interferon-stimulated gene (ISG)-expression screening reveals the specific antibunyaviral activity of ISG20. J. Virol. 92, e02140–17 (2018).

32. Dukhovny, A. et al. A CRISPR activation screen identifies genes that protect against Zika virus infection. J. Virol. 93, e00211–19 (2019).

33. Boys, I. N. et al. RTP4 is a potent IFN-inducible anti-flavivirus effector engaged in a host-virus arms race in bats and other mammals. Cell Host Microbe 28, 712–723.e9 (2020).

34. Catara, G. et al. PARP1-produced poly-ADP-ribose causes the PARP12 translocation to stress granules and impairment of Golgi complex functions. Sci. Rep. 7, 14035 (2017).

35. Dick, A. et al. Role of nucleotide binding and GTPase domain dimerization in dynamin-like myxovirus resistance protein A for GTPase activation and antiviral activity. J. Biol. Chem. 290, 12779–12792 (2015).

36. Fenwick, M. K., Li, Y., Cresswell, P., Modis, Y. & Ealick, S. E. Structural studies of viperin, an antiviral radical SAM enzyme. Proc. Natl. Acad. Sci. U. S. A. 114, 6806–6811 (2017).

37. Gregersen, L. H. et al. MOV10 Is a 5’ to 3’ RNA helicase contributing to UPF1 mRNA target degradation by translocation along 3’ UTRs. Mol. Cell 54, 573–585 (2014).

38. Stokar-Avihail, A. et al. Discovery of phage determinants that confer sensitivity to bacterial immune systems. Cell 186, 1863–1876.e16 (2023).

39. Tan, J. M. J. et al. A DNA-gated molecular guard controls bacterial Hailong anti-phage defence. Nature 643, 794–800 (2025).

40. Zhang, T. et al. Bacterial immune activation via supramolecular assembly with phage triggers. Nature 651, 1051–1059 (2026).

41. Patel, P. H. et al. A pore-forming antiphage defence is activated by oligomeric phage proteins. Nature 651, 1060–1067 (2026).

42. Gizzi, A. S. et al. A naturally occurring antiviral ribonucleotide encoded by the human genome. Nature 558, 610–614 (2018).

43. Yin, Y. W. & Steitz, T. A. Structural basis for the transition from initiation to elongation transcription in T7 RNA polymerase. Science 298, 1387–1395 (2002).

44. Zhao, Y. et al. Neuralized E3 ubiquitin protein ligase 3 is an inducible antiviral effector that inhibits hepatitis C virus assembly by targeting viral E1 glycoprotein. J. Virol. 92, (2018).

45. Abramson, J. et al. Accurate structure prediction of biomolecular interactions with AlphaFold 3. Nature 630, 493–500 (2024).

46. Nishiya, T. et al. Regulation of inducible nitric-oxide synthase by the SPRY domain- and SOCS box-containing proteins. J. Biol. Chem. 286, 9009–9019 (2011).

47. Feng, Y. et al. SPSB1 promotes breast cancer recurrence by potentiating c-MET signaling. Cancer Discov. 4, 790–803 (2014).

48. Liu, S., Nheu, T., Luwor, R., Nicholson, S. E. & Zhu, H.-J. SPSB1, a novel negative regulator of the transforming growth factor-β signaling pathway targeting the type II receptor. J. Biol. Chem. 290, 17894–17908 (2015).

49. Georgana, I. & Maluquer de Motes, C. Cullin-5 adaptor SPSB1 controls NF-κB activation downstream of multiple signaling pathways. Front. Immunol. 10, 3121 (2019).

50. Kato, M., Ito, T., Wagner, G., Richardson, C. C. & Ellenberger, T. Modular architecture of the bacteriophage T7 primase couples RNA primer synthesis to DNA synthesis. Mol. Cell 11, 1349–1360 (2003).

51. Hernandez, A. J., Lee, S.-J., Thompson, N. J., Griffith, J. D. & Richardson, C. C. Residues located in the primase domain of the bacteriophage T7 primase-helicase are essential for loading the hexameric complex onto DNA. J. Biol. Chem. 298, 101996 (2022).

52. Woo, J.-S. et al. Structural and functional insights into the B30.2/SPRY domain. EMBO J. 25, 1353–1363 (2006).

53. Filippakopoulos, P. et al. Structural basis for Par-4 recognition by the SPRY domain- and SOCS box-containing proteins SPSB1, SPSB2, and SPSB4. J. Mol. Biol. 401, 389–402 (2010).

54. Gao, Y. et al. Structures and operating principles of the replisome. Science 363, eaav7003 (2019).

55. Woo, J.-S., Suh, H.-Y., Park, S.-Y. & Oh, B.-H. Structural basis for protein recognition by B30.2/SPRY domains. Mol. Cell 24, 967–976 (2006).

56. Kim, R. S., Levy Karin, E., Mirdita, M., Chikhi, R. & Steinegger, M. BFVD-a large repository of predicted viral protein structures. Nucleic Acids Res. 53, D340–D347 (2025).

57. Cia, G., Kwasigroch, J., Stamatopoulos, B., Rooman, M. & Pucci, F. pyScoMotif: discovery of similar 3D structural motifs across proteins. Bioinform. Adv. 3, vbad158 (2023).

58. Jayaram, H., Taraporewala, Z., Patton, J. T. & Prasad, B. V. V. Rotavirus protein involved in genome replication and packaging exhibits a HIT-like fold. Nature 417, 311–315 (2002).

59. Hughes, F. J. & Romanos, M. A. E1 protein of human papillomavirus is a DNA helicase/ATPase. Nucleic Acids Res. 21, 5817–5823 (1993).

60. Li, T.-F. et al. Human Norovirus NS3 has RNA helicase and chaperoning activities. J. Virol. 92, e01606–17 (2018).

61. Han, K. R. et al. Nucleotide triphosphatase and RNA chaperone activities of murine norovirus NS3. J. Gen. Virol. 99, 1482–1493 (2018).

62. Abbate, E. A., Berger, J. M. & Botchan, M. R. The X-ray structure of the papillomavirus helicase in complex with its molecular matchmaker E2. Genes Dev. 18, 1981–1996 (2004).

63. Zhang, R. et al. A CRISPR screen defines a signal peptide processing pathway required by flaviviruses. Nature 535, 164–168 (2016).

64. Park, R. J. et al. A genome-wide CRISPR screen identifies a restricted set of HIV host dependency factors. Nat. Genet. 49, 193–203 (2017).

65. Scherzinger, E., Lanka, E. & Hillenbrand, G. Role of bacteriophage T7 DNA primase in the initiation of DNA strand synthesis. Nucleic Acids Res. 4, 4151–4163 (1977).

66. Humolli, D. et al. Completing the BASEL phage collection to unlock hidden diversity for systematic exploration of phage-host interactions. PLoS Biol. 23, e3003063 (2025).

67. Zhou, W. et al. Structure of the human cGAS-DNA complex reveals enhanced control of immune surveillance. Cell 174, 300–311.e11 (2018).

68. Studier, F. W. Protein production by auto-induction in high density shaking cultures. Protein Expr. Purif. 41, 207–234 (2005).

69. Langmead, B. & Salzberg, S. L. Fast gapped-read alignment with Bowtie 2. Nat. Methods 9, 357–359 (2012).

70. Danecek, P. et al. Twelve years of SAMtools and BCFtools. Gigascience 10, giab008 (2021).

71. Schuermann, J. P., Perry, K., Neau, D. & Murphy, F. V. RAPD2: Rapid automated processing of macromolecular crystallographic data 2. Struct. Dyn. 12, A280–A280 (2025).

72. Evans, P. R. & Murshudov, G. N. How good are my data and what is the resolution? Acta Crystallogr. D Biol. Crystallogr. 69, 1204–1214 (2013).

73. Liebschner, D. et al. Macromolecular structure determination using X-rays, neutrons and electrons: recent developments in Phenix. Acta Crystallogr. D Struct. Biol. 75, 861–877 (2019).

74. Emsley, P., Lohkamp, B., Scott, W. G. & Cowtan, K. Features and development of coot. Acta Crystallogr. D Biol. Crystallogr. 66, 486–501 (2010).

75. Meng, E. C. et al. UCSF ChimeraX: Tools for structure building and analysis. Protein Sci. 32, e4792 (2023).

76. Zhu, B., Lee, S.-J. & Richardson, C. C. An in trans interaction at the interface of the helicase and primase domains of the hexameric gene 4 protein of bacteriophage T7 modulates their activities. J. Biol. Chem. 284, 23842–23851 (2009).

